# A “loopless” subdomain of transferrin binding protein B can elicit a broadly cross-reactive humoral immune response targeting variants from both encapsulated and non-typeable *Haemophilus influenzae*

**DOI:** 10.64898/2026.09.23.753870

**Authors:** Nikolas F. Ewasechko, David M. Curran, Luisa Samaniego-Barron, Conrad Izydorczyk, Stephen Z. Lee, Michael D. Parkins, Anthony B. Schryvers

## Abstract

*Haemophilus influenzae* is a Gram-negative bacterium that causes pneumonia, otitis media, and invasive infections such as meningitis and bacteremia. A vaccine that confers protection against disease caused by *H. influenzae* serotype b is currently available and is highly effective, but infections caused by non-b serotypes and non-typeable strains are still prevalent and increasing, suggesting a need for a broadly cross-protective vaccine that will protect against infection by all *H. influenzae* strains. To address the need for broad cross-protection, this study focused on the surface lipoprotein component of the bipartite bacterial transferrin receptor, transferrin binding protein B (TbpB), which is universally present *H. influenzae*, irrespective of encapsulation status, and is essential for bacterial colonization and pathogenesis. To prevent vaccine escape, we assessed the sequence and structural diversity among TbpB variants derived from local and international *H. influenzae* isolates and observed that these sequences cluster independently of encapsulation status, suggesting that conferring protection against all *H. influenzae* strains regardless of capsule type or presence of a capsule using a TbpB-based vaccine is feasible. In addition, our diversity analysis also revealed that the C-terminal lobe (C-lobe) of TbpB contains several large, highly variable loops. After immunizing mice with a trivalent vaccine consisting of a representative set of TbpB variants, we found that the resulting antiserum demonstrated modest cross-reactivity against heterologous TbpB variants. However, immunizing with a “loopless C-lobe” (LCL) that lacks the large, highly variable loops we had identified elicited broad cross-reactivity against a panel of intact TbpB variants. This suggests that while the intact TbpB may only generate a moderately cross-reactive antibody response, a vaccine consisting of a single LCL may be able to elicit a sufficiently cross-reactive humoral response against diverse TbpB variants, and in turn, confer broad cross-protection targeting both encapsulated and non-typeable *H. influenzae*.

## INTRODUCTION

*Haemophilus influenzae* is a Gram-negative non-motile coccobacillus that has adapted to survive in the human nasopharynx, its predominant ecological niche (1). Although it can asymptomatically colonize this niche, it is also responsible for a variety of mucosal and invasive infections in humans. Six serotypes, designated a through f, have been identified based on their antigenically distinct polysaccharide capsules, which serve as an important virulence factor, conferring protection against phagocytosis and complement-mediated killing (2). Unencapsulated strains lack this capsule and are thus referred to as non-typeable *H. influenzae* (NTHi). *H. influenzae* serotype b (Hib) is considered the most virulent serotype and has historically been a major etiological agent of invasive infections such as meningitis, epiglottitis, bacteremia, pneumonia, cellulitis, and septic arthritis (3,4). Meanwhile, NTHi strains are more likely to cause mucosal infections such as otitis media, sinusitis, and lower respiratory infections in patients with chronic lung disease, including chronic obstructive pulmonary disease (COPD) – termed “exacerbations” – but are still occasionally responsible for invasive infections (5,6). The advent of a glycoconjugate vaccine targeting Hib in the late 1980s has greatly reduced the incidence of Hib infections; however, it has no effect on NTHi and non-Hib encapsulated strains.

While some progress has been made toward identifying potential vaccine candidates targeting NTHi (7) and *H. influenzae* serotype a (8), the glycoconjugate Hib vaccine remains the only *H. influenzae* vaccine approved for use in humans. For targeting the non-Hib strains, in lieu of producing individual capsular vaccines for each of the six serotypes in addition to a protein-based vaccine for NTHi strains, in our view, a more efficient approach for preventing *H. influenzae* infections would be to target a surface-exposed protein that is essential for survival in the human upper respiratory tract and is thus present in both encapsulated and NTHi strains. Since both encapsulated *H. influenzae* serotypes and NTHi depend on iron acquisition from the iron-sequestering human protein transferrin (Tf) during asymptomatic colonization and invasive infections (9) the Tf receptors expressed on the outer membrane of *H. influenzae* are ideal candidates for use in a broadly cross-protective vaccine targeting both encapsulated and non-encapsulated strains of *H. influenzae*.

In addition to *H. influenzae*, the Tf receptor system is present in other related Gram-negative bacteria that colonize a similar niche in the human upper respiratory tract, such as *Neisseria meningitidis* and *Moraxella catarrhalis* (10,11), as well as veterinary pathogens that colonize the respiratory tissues of food production animals such as pigs and cattle (12,13). In each of these bacteria, the Tf receptor consists of Tf binding protein B (TbpB) and Tf binding protein A (TbpA). TbpB is a surface lipoprotein anchored to the outer membrane that extends into the extracellular milieu, captures iron-loaded Tf, and subsequently brings it in close proximity to the second component of the receptor, TbpA, which is an integral membrane protein that transports iron across the outer membrane after sequestering it away from Tf (14). Although initial attempts to develop a TbpB-based vaccine against meningococcal serogroup B were abandoned in the early phases of a clinical trial (15), new approaches have since been explored. For example, a mutant recombinant TbpB antigen engineered by site-directed mutagenesis such that it was defective in its ability to bind porcine Tf offered superior protection against infection by the porcine pathogen *Glaesserella parasuis* compared to both the intact TbpB and a commercially available inactivated whole-cell vaccine (16). Furthermore, additional findings demonstrated that TbpB mutants defective at binding Tf are capable of eliciting protection against both homologous and heterologous challenge strains (17,18). This suggests that a vaccine consisting of a single TbpB antigen may be capable of conferring broad cross-protection against strains expressing divergent TbpB variants.

Developing vaccines targeting the bacterial Tf receptor is complicated by the natural competence of bacteria such as *H. influenzae*, facilitating the ready uptake of DNA from the extracellular milieu using their efficient natural transformation systems. This allows them to incorporate genetic variations that promote their ability to adapt to selective pressures. The DNA uptake machinery recognizes specific uptake signal sequences (USS) such that they selectively take up DNA from the same or related species (19). To address the issue of potential vaccine escape through antigenic variation, Curran *et al.* (20) conducted a phylogenetic analysis of the sequence diversity among TbpB variants from three important porcine pathogens – *G. parasuis*, *Actinobacillus pleuropneumoniae*, and *Actinobacillus suis* – and found that TbpB sequences derived from these three species distributed into three distinct clades in a manner that showed little correlation with their taxonomic classification. These results point toward a shared genetic reservoir that is accessed by all three species via horizontal gene transfer. It also suggests that a single vaccine composition consisting of a representative TbpB variant from each of the three identified clades may be able to confer protection against infection by all three pathogens – a trend that closely mirrors the conclusions discerned from an analysis of the sequence diversity among meningococcal TbpBs (21). In addition, this study found that the C-terminal lobe (“C-lobe”) of TbpB was much more conserved than the N-terminal lobe (“N-lobe”) and that the antibodies elicited by the C-lobe alone were more broadly cross-reactive against a panel of representative TbpB variants than those elicited by either the intact protein or the N-lobe alone (20). These findings suggested that a TbpB-based vaccine could be strategically designed to confer broad cross-protection by either: a) combining multiple TbpB variants that are representative of each identified phylogenetic cluster in the same vaccine formulation, or b) engineering the protein such that only the most conserved domains of the protein are present in the vaccine. In the present study, we set out to compare these two vaccination strategies. To do this, we first assessed the sequence and structural diversity among *H. influenzae* TbpBs and used the results of this analysis to: i) identify a set of TbpB variants that would be representative of the total antigenic diversity among *H. influenzae* TbpBs, and ii) generate a highly conserved recombinant subdomain of the *H. influenzae* TbpB. We then immunized mice with either a multivalent vaccine containing several representative TbpB variants or a vaccine formulation containing a single conserved TbpB subdomain and compared the cross-reactivity of the humoral response elicited by each TbpB-derived vaccine.

## MATERIALS & METHODS

### Source of *H. influenzae* isolates and mining of reference sequences

The sources for the *H*. *influenzae* isolates used for the sequence diversity analysis component of this study are outlined in Table 1. Strain collection 1 consisted predominantly of clinical isolates either obtained from patients at the Foothills Medical Centre in Calgary, AB, provided by Calgary Lab Services, or donated by Dr. Eric Hansen at the University of Texas Southwestern Medical Center. All strains in collection 1 were stored at -80°C in 16% glycerol, and the original, “un-passaged” stock was used to cultivate bacteria for sequencing whenever possible. The isolates in collection 2 were obtained exclusively from patients with cystic fibrosis (CF) attending the Southern Alberta Adult CF Clinic between 2002 and 2016. The methods used to sample and sequence the genomes of *H. influenzae* isolates from persons with CF are described previously (22) and below. Reference sequences were also obtained from the National Center for Biotechnology Information (NCBI) online database (23) as well as the PubMLST database (24).

**Table 1:** Source of *H. influenzae* TbpB amino acid sequences used in the phylogenetic analysis conducted as part of this study.

| Source | # of TbpB sequences |
| --- | --- |
| Strain collection 1 | 20 (14 Hib, 5 NTHi, 1 *H. influenzae* serotype d (Hid)) |
| Strain collection 2 (CF isolates) | 81 (79 NTHi, 2 Hif) |
| NCBI | 281 |
| PubMLST | 29 |
| Total | 411 |
| Total unique sequences after<br>redundancy removed | **296** (13 Hib, 26 NTHi, 1<br>Hid, 1 Hif, 255 unknown) |

### DNA isolation, amplification, sequencing, and sequence analysis

*H. influenzae* strains from collection 1 were streaked out on chocolate agar plates and the plates subsequently incubated overnight (18-24 h) at 37°C in 5% CO2. The following day, genomic DNA was isolated from the *H. influenzae* cells using the EZNA Bacterial DNA Kit (Omega Biotek). Polymerase chain reaction (PCR) amplification was then used to amplify the *tbpB* genes using primers annealing to sequences located upstream and downstream of each gene (Table 2). Agarose (1%) gel electrophoresis was then used to determine whether the genes were successfully amplified. Following this, the amplicons were purified using the EZNA CyclePure Kit (Omega Biotek) and sequenced by Sanger sequencing, which was performed by the Core DNA Services at the University of Calgary using the same primers used to amplify the *tbpB* genes (Table 2).

**Table 2:**
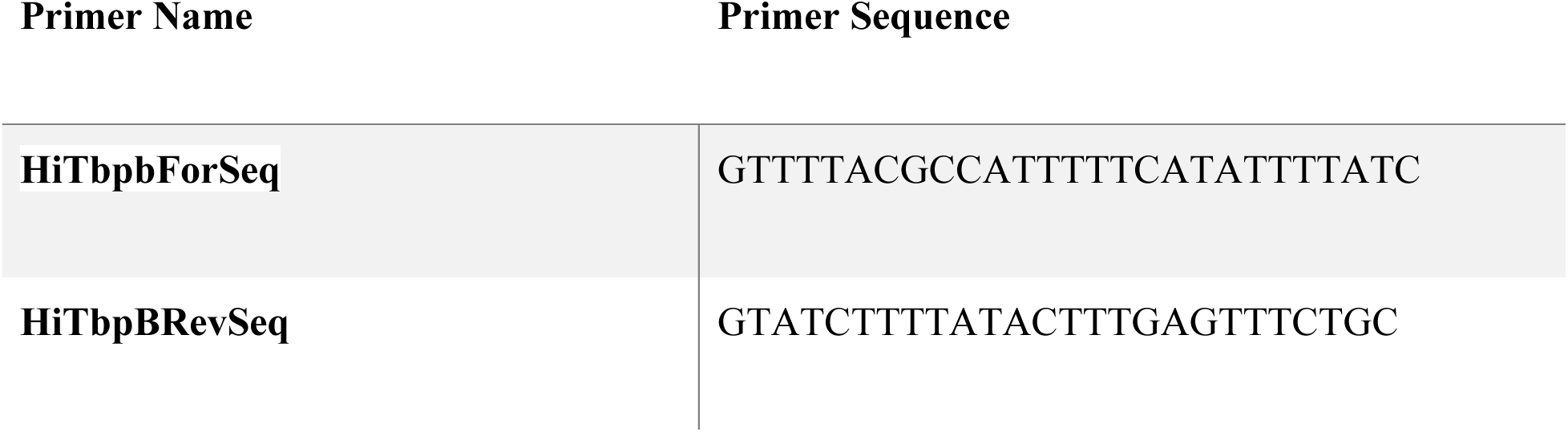
Primers used to amplify and sequence the *tbpB* genes from various strains of *H. influenzae*.

*H. influenzae* isolates in collection 2 were obtained from sputum samples collected from patients with CF attending the Southern Alberta CF Clinic between January 2002 and December 2016. After being identified as *H. influenzae*, each isolate was stored at -80°C in 16% glycerol. Whole-genome sequencing was then performed on a subset of these isolates as described previously (22). To extract the *tbpB* sequences from the sequenced genomes, a nucleotide BLAST database was created for each isolate from its annotated sequences using BLAST+ (v2.11.0) (23). Reference sequences corresponding to *tbpB* were then obtained from *H. influenzae* strain Rd KW20 via NCBI and subsequently used as the query to search each isolate’s database. The top search result for each database was then extracted using a custom Python script.

The resulting TbpB amino acid sequences from all 3 sources (collections 1 and 2 plus online databases) were aligned using Clustal Omega (25), and the aligned sequences were used as input to construct a maximum likelihood tree using RAxML version 8.0 (26) as described in the Supplementary Methods.

### Protein structure modelling and visualization

All structural models depicted in this manuscript were generated using AlphaFold2 (27) and were visualized using PyMOL version 2.5.1 (28).

### Mouse immunizations

Representative TbpBs were first selected using a custom program, “Navargator” (29) (see Supplementary Methods). The three selected *H. influenzae* TbpB antigens were then produced as described previously (16,30) and stored in 100-µL aliquots at -80°C until needed. Immediately prior to immunization, the proteins were thawed and diluted to a concentration of 0.5 mg/mL in Dulbecco’s phosphate-buffered saline (D-PBS, with calcium and magnesium; Multicell). The diluted proteins were then mixed with the squalene-based oil-in-water emulsion adjuvant AddaVax (InvivoGen) such that each vaccine dose contained 50% v/v AddaVax. Forty-eight male C57BL/6 mice (Charles River, 6 weeks old; 8 mice per treatment group) were each immunized with 100 µL of the vaccine formulation, which amounted to a dose of 25 µg of protein. The group receiving the trivalent vaccine, meanwhile, was given 25 µg of each of the three TbpBs (formulated in the same 100-µL dose), and the negative control group received AddaVax alone diluted in D-PBS to a final concentration of 50% v/v. Vaccine doses were administered on days 0, 21, and 42 via the intraperitoneal route. Blood was collected via tail bleed one day prior to the first immunization to obtain “pre-immune” serum. Following the third immunization, the mice were euthanized on day 56, which constituted the experimental endpoint for this study, and a final blood collection was conducted using an intracardiac bleed. Antiserum was then obtained from whole blood by centrifugation (Figure 1).

**Figure 1.**
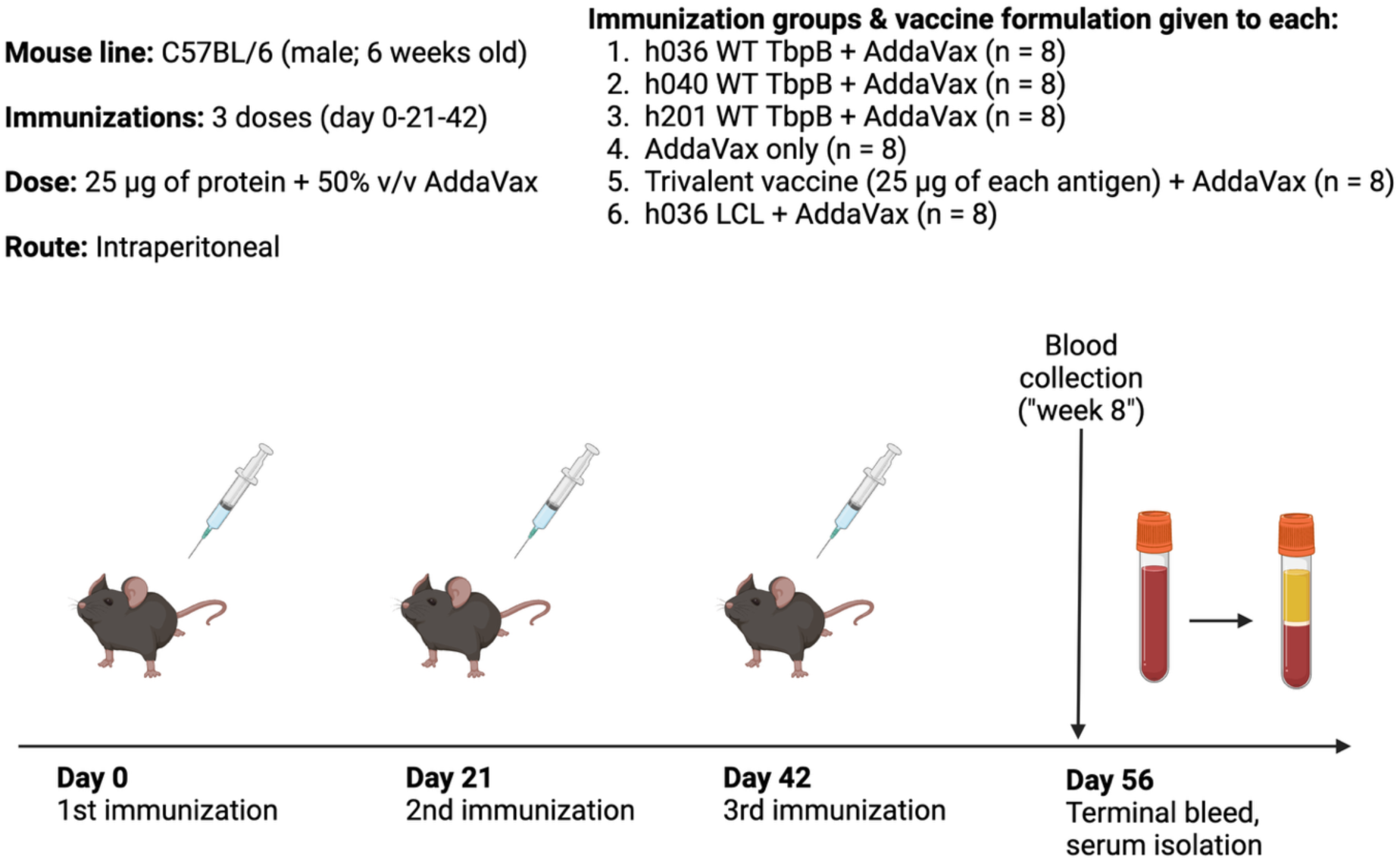
Schematic showing the immunization regimen used in this study. Created using BioRender.com.

### Streptavidin ELISA to detect anti-TbpB IgG titres

To prepare biotinylated protein antigens for coating ELISA plates, recombinant TbpBs were expressed as described previously (16,30), and the resulting *Escherichia coli* ER2566 cell lysates were diluted in PBS + 0.05% Tween-20 (PBST; Sigma-Aldrich) and added to 96-well streptavidin-coated ELISA plates (Greiner BioOne). The lysate-coated plates were then incubated overnight at 4°C. The lysates were removed, and the plates were washed three times with PBST and subsequently blocked using 5% skim milk in PBST for 2 h at room temperature. Following another wash step with PBST, the wells were coated with antiserum diluted in 2.5% skim milk and the plates incubated overnight at 4°C. The next day, the plates were washed and then treated with horseradish peroxidase (HRP)-labelled goat anti-mouse IgG secondary antibody solution (Sigma-Aldrich) diluted by a factor of 10,000 in 2.5% skim milk and the plates incubated for 1 h at room temperature. A final wash step was conducted, and the plates were then developed via the addition of 3,3’,5,5’-tetramethylbenzidine (TMB; Sigma-Aldrich) substrate solution to each well and subsequent incubation for 20 min in a dark compartment at room temperature. The reaction was then quenched by adding 4 N HCl to each well and the resulting OD450 absorbance values determined using an ELISA plate reader (BioTek Synergy HTX Multimode Reader; Agilent). To ensure that the wells on each plate were consistently saturated with properly folded TbpB, several wells per plate were treated with horseradish peroxidase-conjugated human Tf (HRP-hTf) in lieu of the secondary antibody treatment (see Supplementary Figure 1 for a summary of the binding capabilities of all 15 intact TbpB variants used to coat ELISA plates in this study). Additional controls used in the ELISAs conducted for this study included a “no serum” control as well as a “mock lysate” control, which involved coating a row of wells with lysate produced by *E. coli* ER2566 cells in the absence of any plasmid and testing each serum sample against the mock lysate to assess the degree of non-specific binding to endogenous protein antigens expressed by *E. coli*.

### Statistical analyses

Statistical analyses were carried out and figures created using Prism (version 9.0 for macOS; GraphPad). Homologous anti-TbpB IgG endpoint titres and absorbance readings in the cross-reactivity ELISAs were compared using Brown-Forsythe and Welch ANOVA tests followed by Dunnett’s T3 multiple comparisons test. In addition, a two-way ANOVA followed by Šídák’s multiple comparisons test were used to compare fully immunized mice with naïve mice and to compare the cross-reactivity of antiserum generated by the trivalent vaccine formulation with that elicited by the h036 loopless C-lobe (LCL) vaccine antigen.

### Animal welfare

All experiments described in this manuscript involving mice were performed in strict adherence with the experimental protocol approved by the University of Calgary’s Animal Care Committee (protocol #: AC18-0210).

## RESULTS

### Sequence diversity and selection of representatives among TbpB variants

The sequence relationships between the unique *H. influenzae* TbpB variants from the two strain collections, and from the two strain collections plus publicly available databases, are shown in Figures 2A and 2B, respectively. The sequences derived from strain collections 1 and 2 (i.e. the sequences with known serotypes) are interspersed throughout the tree depicted in Figure 2B, suggesting that they are somewhat representative of the total diversity among *H. influenzae* TbpB variants. When looking at either tree in Figure 2, there is no apparent correlation between the TbpB sequence and the serotype of the originating strain, at least for Hib and NTHi; there are too few sequences from serotypes d or f to make an adequate judgement. This pattern is often observed in organisms that exhibit high rates of horizontal gene transfer and is a part of our rationale for developing a capsule-agnostic vaccine. The sequences in the small tree fall into a few major clades, though as the branch lengths between clades are much smaller than the branch lengths within each clade, such a classification is not particularly rigorous or well-resolved. We used Navargator to select 3 representative variants from this tree, which yielded: h031, h038, and h040. However, because of the availability of expression vectors encoding the h036 and h201 variants prior to conducting this analysis, these were chosen to replace h031 and h038, respectively. These replacement pairs share 99% sequence identity, and their selection only decreased the overall clustering score by 0.1%, indicating the two choices should be functionally equivalent. These 3 sequences are referred to as the “immunization panel” (Figure 2A; Table 3).

**Figure 2.**
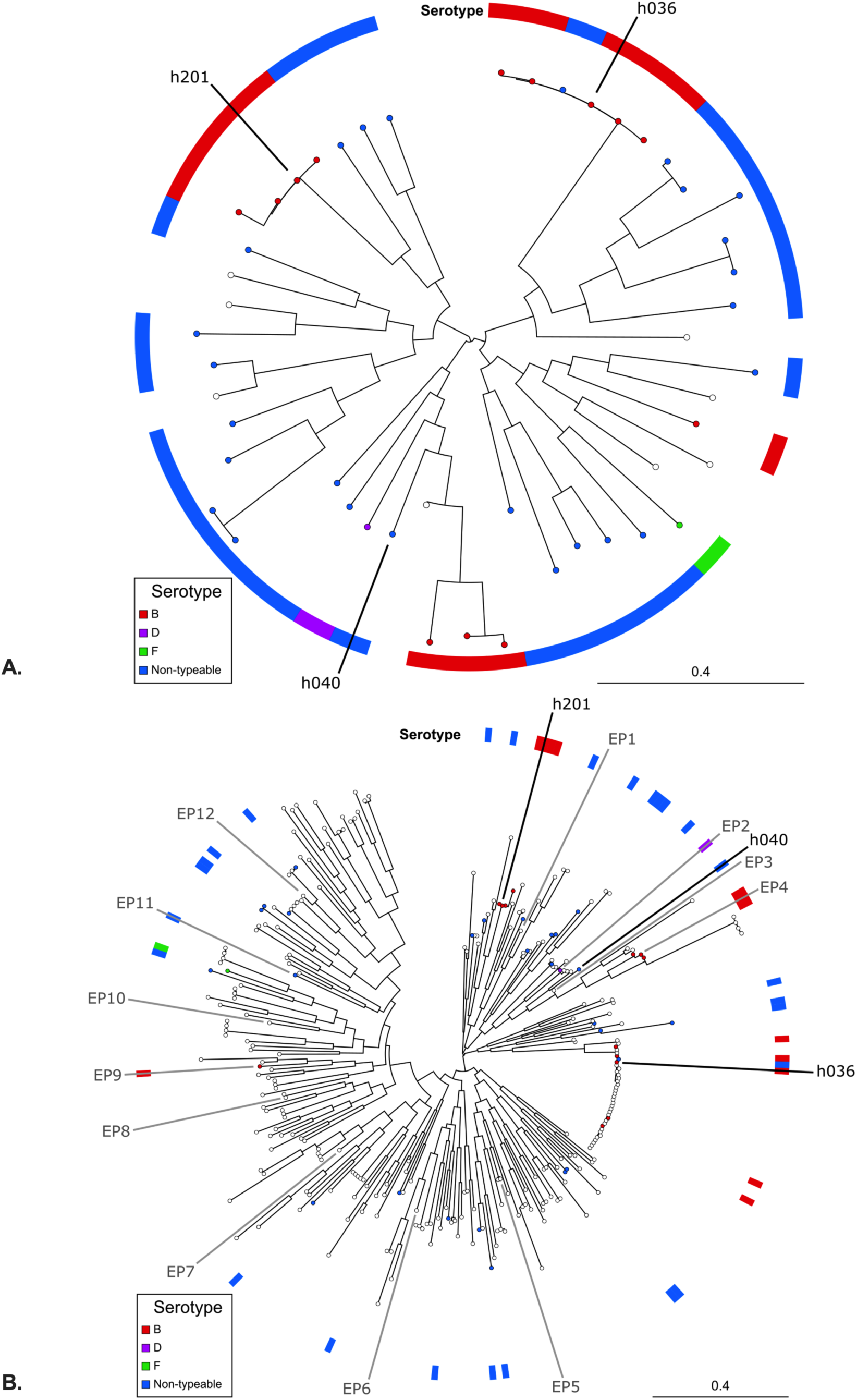
Phylogenetic trees depicting the sequence diversity among *H. influenzae* TbpB variants. (**A**) shows 49 variants from strain collections 1 and 2, while (**B**) shows 296 variants after the addition of unique publicly available sequences. In both trees the serotype of the originating strain, where available, is indicated by the colour of the tree node and by the coloured ring banner around the tree. The locations of the 3 immunizing antigens – h036, h040, and h201 – are indicated in both trees. (**B**) also shows the locations of the 12 additional members of the ELISA panel, labeled TbpB-1 through TbpB-12. Trees were visualized and annotated using Navargator.

**Table 3:** *H. influenzae* TbpB variants used for immunizations (“Immunization panel”) and to coat ELISA plates used in assessing the cross-reactivity of the antiserum elicited by the immunogens (“ELISA panel”). The 3 variants in the immunization panel were also used in the ELISA panel in order to determine the reactivity of the antisera to the homologous variant. The serotype of each strain from which the TbpBs were derived, if known, is also listed beside each variant.

| Immunization Panel | ELISA Panel | Accession | Serotype |
| --- | --- | --- | --- |
| h036 | h036 | h036 | b |
| h040 | h040 | h040 | NT |
| h201 | h201 | h201 | b |
|  | TbpB-1 | WP_050848381.1 |  |
|  | TbpB-2 | h014 | d |
|  | TbpB-3 | WP_112083867.1 |  |
|  | TbpB-4 | h026 | b |
|  | TbpB-5 | WP_221267536.1 |  |
|  | TbpB-6 | WP_118806425.1 |  |
|  | TbpB-7 | PRI69814.1 |  |
|  | TbpB-8 | PRI75995.1 |  |
|  | TbpB-9 | h216 | b |
|  | TbpB-10 | WP_111689186.1 |  |
|  | TbpB-11 | h210 | NT |
|  | TbpB-12 | WP_118872958.1 |  |

As with the smaller tree, our tree of the total unique diversity among *H. influenzae* TbpBs (Figure 2B) depicts a few major clades that are not particularly well-resolved. While our immunization panel is well spread out around the smaller tree, the three sequences are all found in one region of the larger tree that accounts for a little more than 1/3 of the total variants. While this could mean that a final formulation of our vaccine against *H. influenzae* may also include an antigen from the other side of the tree, in this study it provides an opportunity to evaluate how cross-reactivity behaves across a wide range of phylogenetic distances. In order to evaluate this cross-reactivity, we used Navargator to select 10 representative variants from the large tree. This set of 10 variants included h201 as well as a variant that was 99% identical to and subsequently replaced by h036, both from our immunization panel. We added h040, the final member of the immunization panel, as well as 4 variants from strain collection 1 that had previously been used as approximate representatives prior to the availability of Navargator. This gave us a total of 15 variants – the three from the immunization panel plus twelve new variants – that we refer to as the ELISA panel (Table 3).

### Structural diversity among TbpB variants

A Clustal Omega alignment of representative TbpB variants was conducted and revealed that much of the sequence diversity is localized in the N-lobe – specifically, the human Tf (hTf)-binding interface – and in loops 17, 23, and 31 of the C-lobe (see Figure 3 for a schematic depicting the secondary structural elements present in the *H. influenzae* TbpB and Figure 4 for the structural diversity analysis). A model of the structure of the h036 TbpB was then generated using AlphaFold2 (Figure 4B and 5A) and the consensus sequence identity depicted in the alignment mapped onto the h036 TbpB model (Figures 4B and 4C). AlphaFold2 models of the protein structures of the h040 and h201 TbpB variants were also generated (Figures 5C and 5D), which allowed for a side-by-side comparison to be carried out among the 3 intact TbpB immunogens used in this study. This comparison served to highlight the significant size discrepancy among certain loops in the C-lobe – in particular, loop 23, which varied in size between 11 (h040 TbpB) and 37 (h201 TbpB) amino acids (Figure 5). Taken together, the significant size and sequence variability in loop 23 among different TbpB variants and the substantial sequence diversity in the N-lobe as well as loops 17 and 31 appeared to suggest that removing these highly variable regions could result in a broadly cross-protective vaccine antigen, which is what prompted the design and production of the LCL depicted in Figure 5B (see also: Supplementary Figure 2).

**Figure 3.**
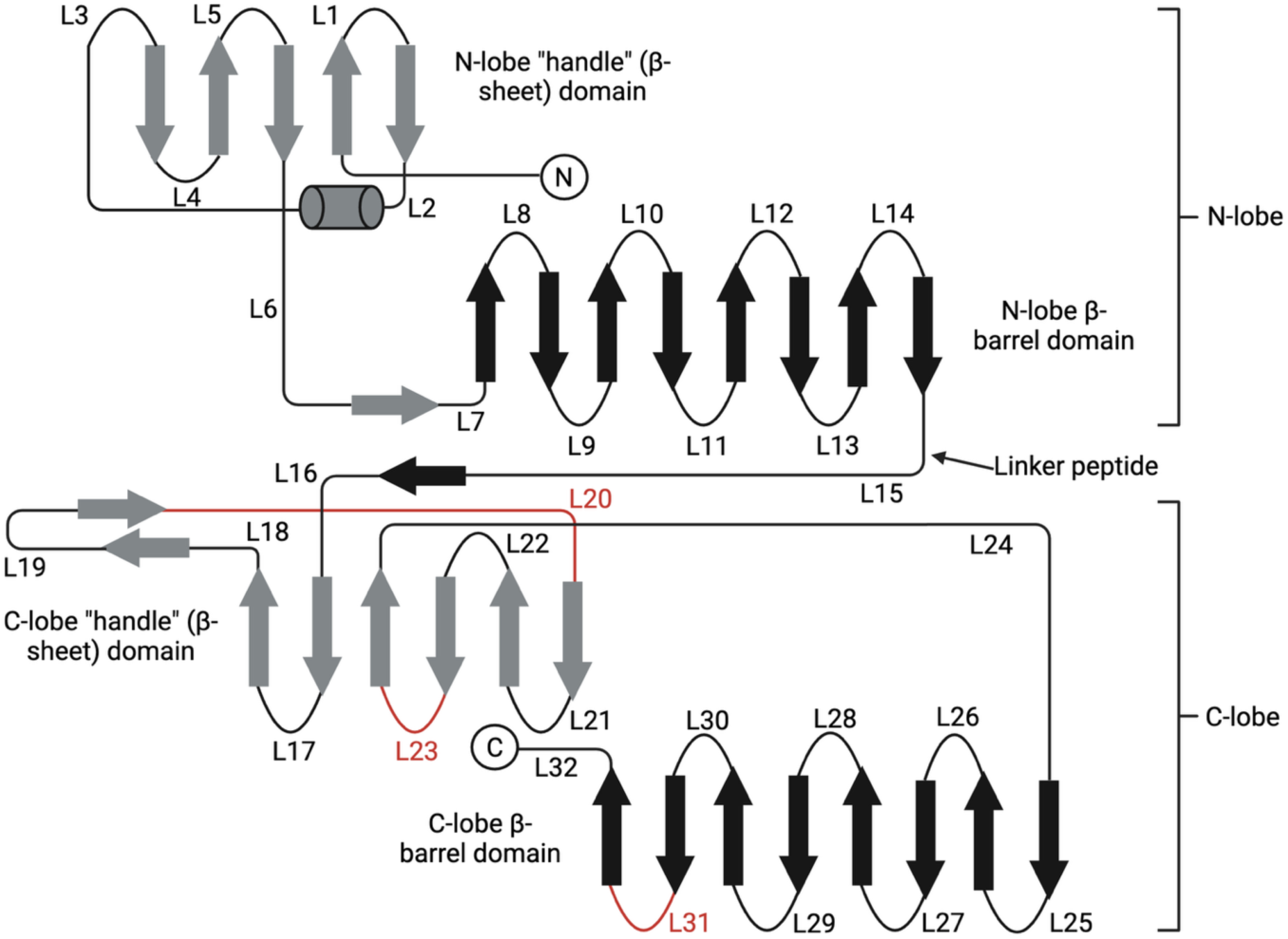
Schematic depicting the principal secondary structural elements present in the *H. influenzae* TbpB. The loops modified in the creation of the LCL used to immunize mice in this study are highlighted in red. Created using BioRender.com.

**Figure 4.**
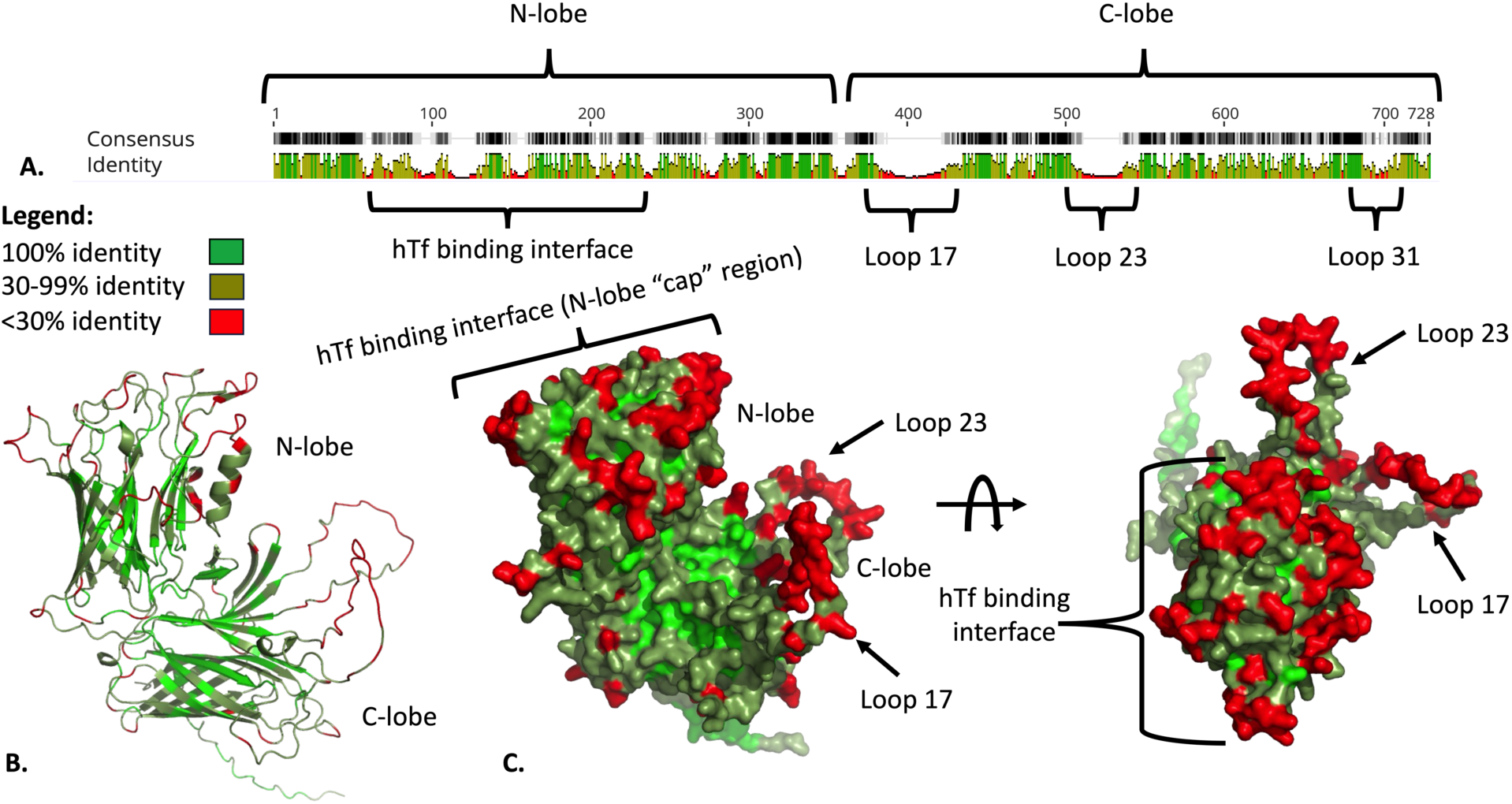
Patterns of sequence and structural diversity among *H. influenzae* TbpBs. (**A**) depicts the “consensus identity” (defined as the mean pairwise identity among all pairs in the column) among 296 unique *H. influenzae* sequences based on an alignment of these sequences carried out using Clustal Omega (sliding window size: 1). The resulting alignment was then visualized using Geneious. (**B**) and (**C**) show the consensus identity depicted in (**A**) mapped onto an AlphaFold2 model of the h036 TbpB. The model was visualized using PyMOL and manually colour-coded according to the above legend.

**Figure 5.**
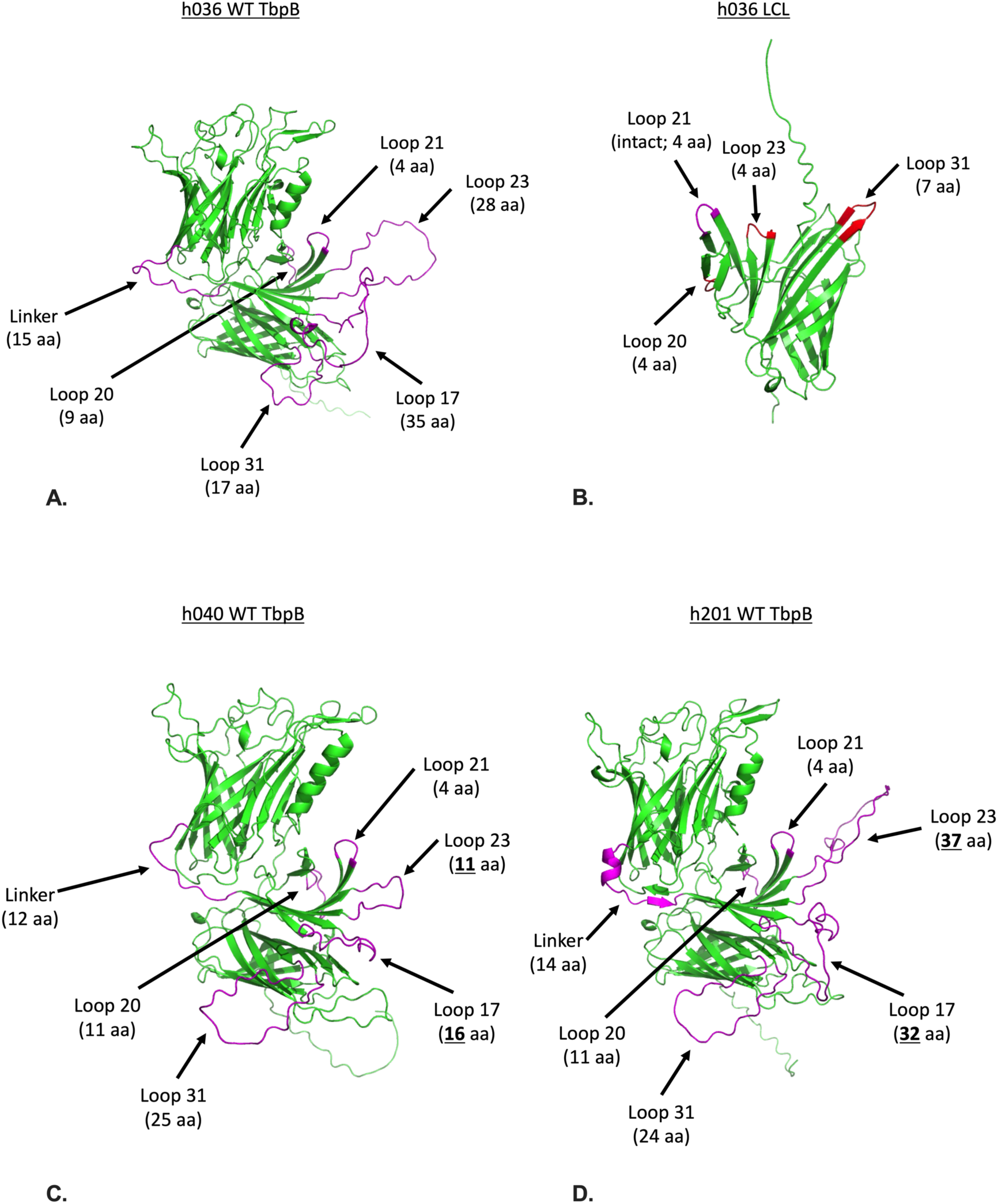
AlphaFold2 models of the four TbpB antigens used to immunize mice in this study. Loops that have been found to contain a substantial amount of variability in size and sequence among different TbpB variants are highlighted in magenta and the size of each loop specified beneath each label. In (**B**), the loops that were replaced with smaller loops from the *A. pleuropneumoniae* TbpB (loops 20, 23, and 31) are labelled in red. The endogenous loop 21 was left intact due to it being the same size as the loop that would have replaced it.

### Immunogenicity and cross-reactivity of TbpB-based vaccine formulations

Following the immunization of mice with three representative TbpB variants – formulated individually as well as together in a trivalent vaccine – and one LCL derived from the h036 TbpB, serum samples were collected and the homologous anti-TbpB IgG titres corresponding to each sample were detected by ELISA. All four antigens elicited a robust humoral response, with the h036 wild-type (WT) TbpB appearing to be the most immunogenic (Figures 6A and 6D). In general, the IgG titres elicited by the individually formulated, “monovalent” vaccines were not significantly higher than those elicited by the trivalent vaccine formulation (Figures 6A-C), suggesting that combining the three TbpB variants in the same vaccine formulation likely does not diminish the anti-TbpB humoral response generated by each individual variant. In fact, the trivalent vaccine appeared to produce higher levels of anti-h036 TbpB IgG than both the monovalent h036 TbpB vaccine and the h036 LCL (Figure 6A).

**Figure 6.**
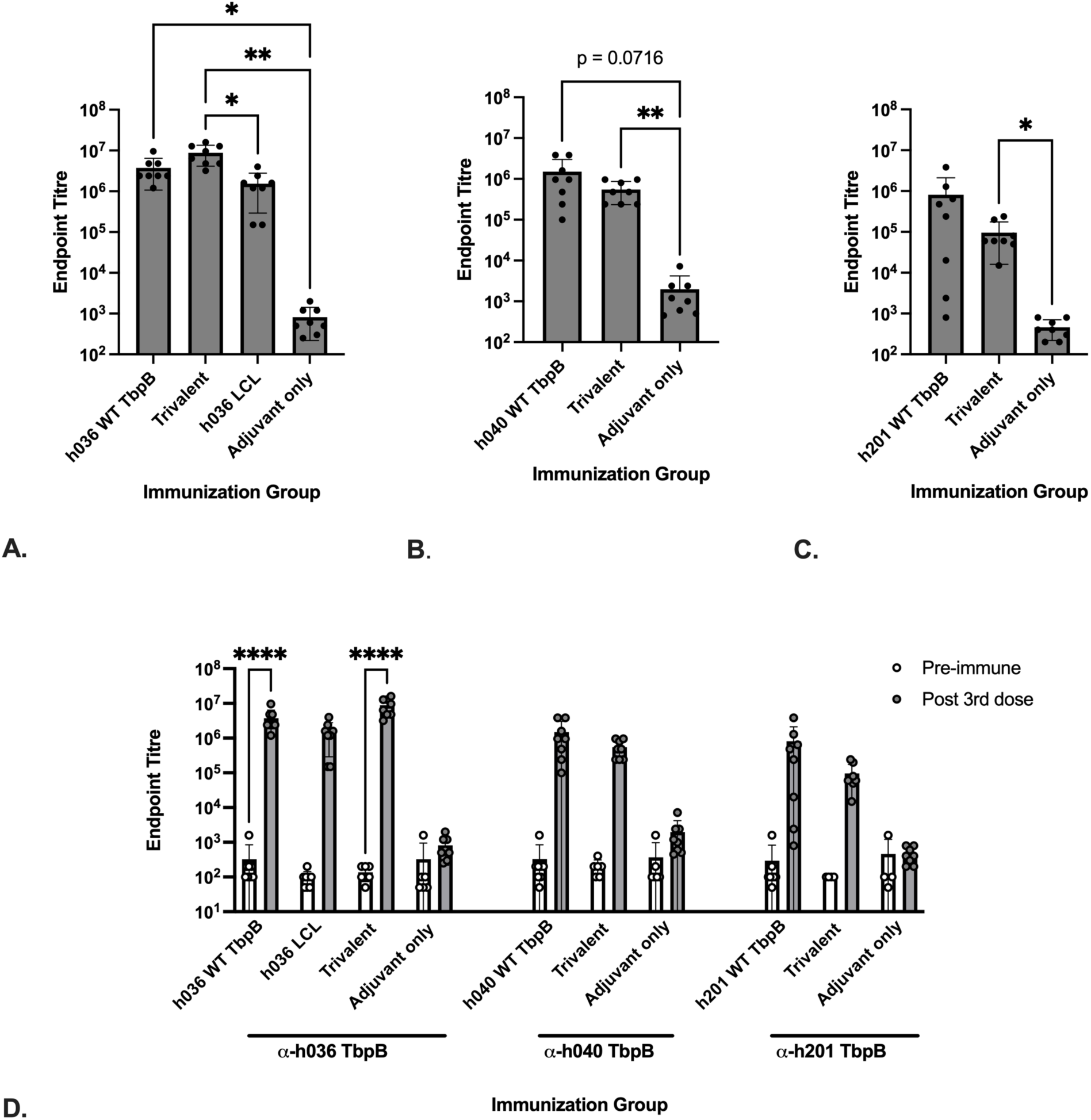
Anti-TbpB IgG titres targeting the three variants in the immunization panel – h036 (**A**), h040 (**B**), and h201 (**C**) – detected in serum samples collected during week 8 from mice immunized with an intact TbpB individually, the trivalent vaccine, the h036 LCL, or adjuvant alone. (**D**) depicts a side-by-side comparison of the mean endpoint titres of serum samples collected post 3^rd^ dose during week 8 to those of the pre-immune serum samples. Samples collected from mice immunized with individual TbpBs were assessed for their reactivity against the respective homologous variant while samples collected from the trivalent and “adjuvant only” groups were tested against all three variants. The h036 LCL, meanwhile, was assessed for its reactivity against the intact h036 TbpB (**A**). Endpoint titres were defined as the reciprocal of the last dilution in the series that produced an absorbance value that was greater than double the negative (no serum) control. ELISA plates used to detect the IgG titres depicted here were coated with lysates containing recombinant TbpB in the absence of MBP. Significance was determined using the Brown-Forsythe and Welch ANOVA tests, followed by a Dunnett’s T3 multiple comparisons test comparing each group to one another (*p<0.05, **p<0.01) (**A**-**C**). In (**D**), mean IgG titres in naïve mice and fully immunized mice samples were compared using a two-way ANOVA followed by Šídák’s multiple comparisons test to determine significance (****p<0.0001). Bars represent the mean (±SD) endpoint titre for sera collected from 8 mice. Two technical replicates were performed, and the data plotted here represent the average endpoint titre detected in each sample in the two replicate experiments.

To ascertain the cross-protective potential of the TbpB-based vaccines used in this study, the cross-reactivity of antiserum elicited by the LCL and the three intact TbpB variants in the immunization panel was tested using ELISA plates coated with a 15-variant panel of representative *H. influenzae* TbpBs. The results of these ELISAs showed that the breadth of cross-reactivity of the antisera elicited by the trivalent vaccine against heterologous variants in the ELISA panel was minimal (Figures 7A and 7G) whereas the sera elicited by the LCL was broadly cross-reactive against the full panel (Figures 7B and 7G). It is interesting to note that, other than the LCL and except for the trivalent sera reacting to TbpB-7, every single significant cross-reactive interaction was against TbpB-1 through TbpB-4 (Figure 7), which are the members of the ELISA panel from the same major region of the tree as the immunization panel (Figure 2B). These included sera elicited by the h036 WT TbpB, which cross-reacted strongly against TbpB-3 and TbpB-4 (Figure 7C), as well as antibodies generated by the h040 WT TbpB, which recognized Tbp-3 (Figure 7D).

**Figure 7.**
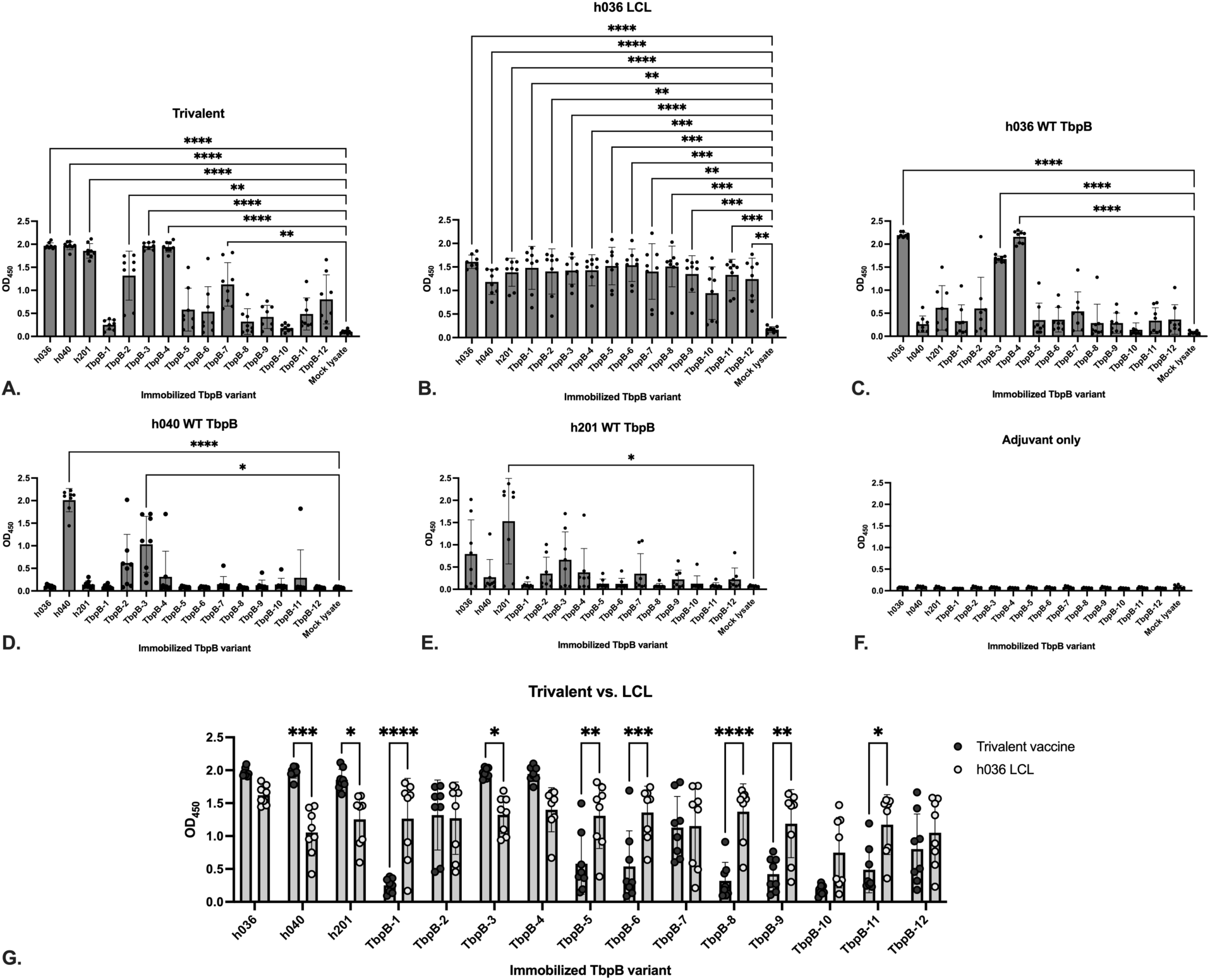
Cross-reactivity of the anti-TbpB antibody response elicited in mice immunized with the trivalent vaccine (**A**), the h036 LCL (**B**) or individually with intact TbpB variants (**C**-**F**) against the ELISA panel developed for this study. (**G**) depicts a direct comparison of the cross-reactivity of the antiserum elicited by the trivalent vaccine and the LCL. A serum dilution of 1/1,000 was used for all samples assessed in this ELISA, and the plotted values represent the raw absorbance readings at 450 nm (OD_450_) produced by each serum sample. Significance was determined using the Brown-Forsythe and Welch ANOVA tests, followed by a Dunnett’s T3 multiple comparisons test comparing each mean to the “mock lysate” mean (*p<0.05, **p<0.01, ***p<0.001, ****p<0.0001). In (**G**), OD_450_ values produced by the trivalent and LCL groups were compared using a two-way ANOVA followed by Šídák’s multiple comparisons test to determine significance (*p<0.05, **p<0.01, ***p<0.001, ****p<0.0001). Bars represent the mean (±SD) absorbance value for sera collected from 8 mice. Two technical replicates were performed, and the data plotted here represent the average absorbance value produced by each sample in the two replicate experiments.

## DISCUSSION

In this study, the sequence and structural diversity among *H. influenzae* TbpBs was assessed, and the results of this analysis was used to inform the strategic selection and design of recombinant TbpB-based vaccine antigens. The sequence diversity analysis revealed that there is little correlation between the encapsulation status of the isolates from which the TbpB sequences used in this analysis were obtained and the clustering patterns in the resulting trees (Figure 2). This suggests that the genetic reservoir contributing to TbpB sequence variation is likely shared amongst both encapsulated and non-typeable *H. influenzae* – a finding that closely aligns with previous reports assessing the sequence diversity among TbpB variants in porcine pathogens (20) and supports the notion that protection against infection by both encapsulated *H. influenzae* strains and NTHi can be conferred with a single TbpB-based vaccine. In addition, TbpB sequence alignments (Figure 4) and structural models of representative TbpB variants (Figure 5) revealed significant variability in the N-lobe as well as specific loops in the C-lobe of the protein. These preliminary observations regarding the patterns of sequence and structural diversity among *H. influenzae* TbpBs ultimately prompted the use of two distinct vaccination strategies, with the overall aim of maximizing the cross-protective potential of TbpB-based vaccines targeting *H. influenzae*: i) combining three representative intact TbpB variants in a single, trivalent vaccine formulation, and ii) generating an LCL that lacked the loops in the C-lobe that featured significant size and sequence variability.

Mice were immunized with each of the two formulations, with the results clearly demonstrating the superior cross-reactivity of an LCL-based vaccine to that of an intact TbpB vaccine (Figures 7A, 7B, and 7G), even when a set of three TbpB variants that, taken together, are somewhat representative of the sequence diversity among *H. influenzae* TbpBs are included in the same vaccine formulation. These results align with previous findings suggesting that a vaccine containing a single TbpB C-lobe derived from the porcine pathogen *A. pleuropneumoniae* elicits antiserum with broadly cross-reactive properties (20). Interestingly, however, they contradict the more recent finding that an intact “cluster III” TbpB derived from another porcine pathogen, *G. parasuis,* confers 100% protection against infection by a *G. parasuis* strain expressing a TbpB from an entirely different cluster, “cluster I” (18). In contrast to the intact *G. parasuis* TbpB, the limited cross-reactivity of a vaccine containing intact *H. influenzae* TbpBs observed in this study appears to align with the results of a previous report that showed that, despite finding that immunizing with a recombinant TbpB-based vaccine promoted clearance of the homologous NTHi strain from the lung in a rat infection model, the antisera elicited by the TbpB antigen exhibited limited bactericidal activity against heterologous strains (31). The underlying mechanisms as to why intact *H. influenzae* TbpBs, unlike TbpBs from porcine pathogens, seem to elicit an antibody response that is minimally cross-reactive against heterologous TbpB variants have yet to be fully elucidated. However, in general, the C-lobe loops of porcine pathogen TbpBs are significantly smaller than those of the *H. influenzae* TbpBs examined in this study (16,32,33), which may be a contributing factor toward the increased cross-reactive and cross-protective properties of *A. pleuropneumoniae* and *G. parasuis* TbpBs compared to intact *H. influenzae* TbpB variants. Hence, it may be that the presence of large, highly variable loops in the *H. influenzae* TbpB C-lobe is restricting the cross-reactivity of the antibody response elicited by these antigens. Indeed, given that the *H. influenzae* LCL antigen used in this study consists of a TbpB C-lobe with substantially reduced loops 23 and 31 (Figure 5B), a reasonable hypothesis might be that loops 23 and 31 are immunodominant and that the presence of these loops diverts the immune response away from more conserved regions of the protein as well as regions that are affected by functional constraints such as the binding of hTf by the N-lobe cap region (Figure 4).

The broad cross-reactivity of the *H. influenzae* LCL highlights the merits of a phylogenetics- and protein structure-based vaccine design approach. The inclusion of multiple representative variants in a single vaccine formulation yielded a less cross-reactive vaccine than the LCL approach in this study; however, our analysis of the sequence diversity among *H. influenzae* TbpB variants combined with the availability of advanced structural modelling tools such as AlphaFold2 (27) allowed us to map the sequence diversity onto structural models of TbpB. In turn, this allowed us to generate a conserved subdomain that would be expected to elicit a cross-reactive antibody response. While the *H. influenzae* TbpB appears to have structural idiosyncrasies – namely, large, highly variable loops – that may preclude the ability of the intact protein to elicit broad cross-protection, approaches using phylogenetics and structural prediction have been used successfully in the past in the selection of representative variants to include in multivalent protein-based vaccine. A notable recent example of this was the use of phylogenetic analyses to select representative factor H binding protein (fHbp) variants to include in a bivalent meningococcal vaccine (Bivalent rLP2086, also known as “Trumenba,” Pfizer), which served to expand the breadth of cross-protection conferred against divergent *N. meningitidis* serogroup B strains (34). A similar approach is also currently being used to develop and evaluate a bivalent TbpB vaccine targeting *Neisseria gonorrhoeae*, yielding promising preclinical results thus far (35). Nevertheless, while the LCL represents a promising vaccine antigen design approach that could feasibly be used to target both encapsulated and non-typeable *H. influenzae* with a single vaccine composition, significant challenges inherent to the development of an NTHi vaccine remain. These pitfalls were highlighted by the recent development and evaluation of a protein-based vaccine targeting surface antigens from both NTHi and *M. catarrhalis* (Mcat) that was designed to prevent exacerbations of COPD. While early clinical trial results for the NTHi-Mcat vaccine (36), including a four-year follow-up study (7), demonstrated a good reactogenicity and immunogenicity profile, the vaccine was later found to provide no discernible benefit toward reducing the yearly rate of COPD exacerbations in patients with COPD who received the vaccine compared to the placebo group (37). Similarly, a phase 2 trial evaluating an iteration of the NTHi vaccine lacking the Mcat antigen UspA2 showed a modest efficacy rate that was ultimately not statistically significant (38). The reasons as to why the primary endpoint of these trials was not met have yet to be elucidated; however, it may be related to the complex interplay between bacterial colonization and the onset of COPD exacerbations, which may require new correlates of protection to be devised for this unique patient population. In any case, it is clear that when evaluating novel NTHi vaccine candidates, both the antigen candidate and the preclinical infection models need to be carefully selected to reflect the natural history of the disease state the vaccine is aiming to prevent.

There are several limitations pertaining to the results presented in this paper that merit further discussion. The first is that there are inherent limitations in deriving structural and functional insights from computational models of protein structures alone. For definitive conclusions regarding the structural features of the *H. influenzae* TbpB to be drawn, the structures of representative TbpB variants and/or TbpB subdomains such as the LCL would have to be determined using techniques such as X-ray crystallography or cryo-electron microscopy. Secondly, while the ELISA data depicting the cross-reactivity of IgG molecules against a diverse panel of *H. influenzae* TbpB variants offer a promising path forward toward the development of a broadly protective protein-based vaccine targeting both encapsulated *H. influenzae* and NTHi, the h036 LCL has yet to be evaluated for its ability to confer protection against infection *in vivo* using a suitable animal infection model. Unfortunately, this is an endeavour that is complicated by the need to supplement *H. influenzae* with “virulence enhancement agents” in order to induce invasive disease in rodent infection models (39). Nevertheless, efforts are ongoing to leverage the robust meningococcal infection model with which our group has extensive experience (40,41) by engineering meningococcal strains demonstrated to be virulent in mice when supplemented with human Tf such that they express the *H. influenzae* Tf receptor. Additionally, as this study focused solely on IgG titres, additional antibody subtypes such as IgA will also need to be assessed in future experiments for their cross-reactive properties against divergent *H. influenzae* TbpBs. Moreover, the immunizations conducted as part of this study should be repeated using female mice to ensure that the cross-reactivity trends observed herein using serum derived from male mice are also observed when serum taken from female mice is assessed.

Overall, the present study demonstrates the utility of sequence and structural diversity analyses to circumvent vaccine escape caused by a significant amount of antigenic variability in a vaccine target. Given their essential function for host colonization and niche adaptation (42), Tf receptors are under constant selection pressure exerted by the host immune system, resulting in the emergence of diverse TbpB variants through horizontal gene transfer among related colonizing bacteria (19). While conventional wisdom would suggest that a highly variable antigen be avoided when it comes to the selection of vaccine candidates, in this paper, we show that TbpBs can be strategically engineered such that only the most conserved regions of the protein are present and that this conserved subdomain can elicit a broadly cross-reactive humoral immune response. Thus, the h036 LCL represents a promising protein-based vaccine candidate that, due to the ubiquity of Tf receptors across all *H. influenzae* strains (9), has the potential to confer protection against all non-Hib *H. influenzae* infections, which remain a significant and underappreciated global health burden (43).

### Ethics Statement

The mouse immunization studies were conducted in accordance with animal use protocol AC18-0210, which was reviewed and approved by the Animal Care Committee at the University of Calgary’s Health Sciences Animal Resource Centre.

## Acknowledgements

We would like to thank Dr. Jamie Fegan (U. of Toronto) for her advice on how best to present the immunological data collected as part of this study and for providing insights/feedback throughout the conceptualization and data collection phases of this study.

