## Supplementary Methods for "A “loopless” subdomain of transferrin binding protein B can elicit a broadly cross-reactive humoral immune response targeting variants from both encapsulated and non-typeable *Haemophilus influenzae*"

### **Sequence alignment and phylogenetic analysis:**

The TbpB amino acid sequences derived from the *H. influenzae* strain collections 1 and 2 were combined with reference sequences downloaded from the NCBI and PubMLST databases. The signal peptide sequences were then removed using SignalP 6.0 (1), and any duplicate sequences were removed using the “Remove Redundancy” feature in Jalview (2). The resulting sequences were aligned using Clustal Omega (3). Subsequently, the aligned sequences were used as input to construct a maximum likelihood tree using RAxML version 8.0 (4). The following command was used to generate the tree:

./raxmlHPC-PTHREADS-AVX2 -f a -p 00001 -m PROTGAMMAWAG -s [input file name] -x 00001 -# 200 -T 10 -n [output file name]

The resulting trees were visualized and annotated using FigTree version 1.4.4 (http://tree.bio.ed.ac.uk/software/figtree/). In addition to being used to generate maximum likelihood trees, the alignment produced by Clustal Omega was also visualized and annotated using Geneious version 2023.2.1 ([https://www.geneious.com](http://www.geneious.com/)).

### **Selection of representative *H. influenzae* TbpB variants:**

Representative TbpB variants were selected using the custom phylogenetics software Navargator (5), which was created to perform a clustering analysis on a phylogenetic tree in order to select variants at the centre of each cluster. The Newick (.nwk) file corresponding to the maximum likelihood tree depicting the sequence diversity among TbpB variants in strain collections 1 and 2 was used as input to the program, with the “Variants to find” parameter set to 3. The 3 variants selected for mouse immunizations will be referred to henceforth as the “immunization panel.” To assess the cross-reactivity of the anti-TbpB sera elicited by our immunization panel, we used Navargator to select 10 representative variants from the large tree consisting of 296 unique sequences. These 10 variants were combined with 5 additional variants as described in the Results section and will be referred to as the “ELISA panel.” The variants belonging to the immunization and ELISA panels are summarized in Table 3.

### **Production and purification of recombinant TbpBs:**

To prepare protein antigens for mouse immunizations, genes encoding *H. influenzae* TbpB variants were first cloned into a custom T7 expression vector (Addgene #: pE5770; Supplementary Figure 3) using restriction cloning (see Supplementary Tables 1 and 2 for additional information). The custom vector encoded an N-terminal maltose binding protein (MBP) fusion partner as well as a polyhistidine tag upstream of the MBP fusion partner to facilitate purification of the protein by affinity chromatography and a tobacco etch virus (TEV) protease cleavage site downstream of the fusion partner to facilitate its removal. This vector was then transformed into chemically competent *Escherichia coli* ER2566 cells. Following a 1-h post-heat shock recovery period at 37°C (with shaking), 6 mL of lysogeny broth (LB) medium containing 100 µg/mL of ampicillin was added and the resulting “starter” culture incubated at 37°C with shaking for 4 h. Next, 6 mL of the starter culture was then used to inoculate 6 L of ZYP-5052 “autoinduction” medium (6) containing 50 µg/mL of ampicillin. The resulting culture was then incubated at 37°C for 18 h with shaking, followed by an incubation at 20°C for 24 h. Following this, the cells were harvested by centrifugation at 5,000 x g for 25 min at 4°C and the cell pellets subsequently re-suspended in 150 mL of Resuspension Buffer (50 mM Tris pH 8.0, 300 mM NaCl, 10 mM imidazole) containing a protease inhibitor tablet (Roche), lysozyme (final concentration: 0.4 mg/mL; Sigma-Aldrich), and DNase I (final concentration: 6.67 µg/mL; Sigma-Aldrich). The cells were then lysed using a high-pressure cell homogenizer (Avestin Emulsiflex C3 Homogenizer). Next, the lysate was centrifuged at 35,000 x g for 90 min at 4°C and the resulting supernatant syringe-filtered (0.2-µm, Pall) and loaded onto the 5-mL HisTrap High Performance Ni-NTA column (Cytiva) in a closed-circuit loop for affinity capture. Following overnight circulation of the clarified lysate through the Ni-NTA column, the column was washed with Wash Buffer (50 mM Tris pH 8.0, 1 M NaCl, 20 mM imidazole) and the MBP-TbpB fusion protein eluted using Elution Buffer (50 mM Tris pH 8.0, 300 mM NaCl, 300 mM imidazole). In preparation for cleavage of the fusion protein using TEV protease, the eluted protein sample was then transferred to dialysis tubing and dialyzed overnight at 4°C against Exchange Buffer (50 mM Tris pH 8.0, 600 mM NaCl). Following overnight dialysis, TEV protease (prepared in-house; 100 µg of TEV protease added per mg of protein) was then added to the protein sample and the sample incubated overnight at 4°C to separate the TbpB from its MBP fusion partner in preparation for further purification.

Following cleavage of the fusion protein by TEV protease, the TbpB was dialyzed against Equilibrium Buffer (50 mM Tris pH 8.0, 10 mM NaCl) overnight at 4°C. The protein was then concentrated to a volume of 10 mL using a Vivaspin 20 concentrator with a 50-kDa molecular weight cut-off (Cytiva) and subsequently loaded onto a HiTrap Q Sepharose High Performance (QHP) column (Cytiva) for anion-exchange chromatography. The column was washed with Equilibrium Buffer and any bound proteins eluted using QHP Elution Buffer (50 mM Tris pH 8.0, 2 M NaCl). As this purification step was carried out using the NGC Chromatography fast-purification liquid chromatography (FPLC) system (BioRad), the UV absorbance (280 nm) readings in the chromatogram produced by the FPLC system were used to detect the presence of protein in each eluted fraction. The fractions with high absorbance were then assessed by sodium dodecyl sulfate-polyacrylamide gel electrophoresis (SDS-PAGE) in order to determine whether TbpB is present in the fraction (post-purification SDS-PAGE gels depicting each of the immunogens used in this study can be seen in Supplementary Figure 4). The fractions that were determined to contain TbpB were then exchanged into phosphate-buffered saline (PBS; 137 mM NaCl, 2.7 mM KCl, 8 mM Na_2_HPO_4_, 2 mM KH_2_PO_4_), concentrated down to between 1 and 10 mg/mL, divided into 100- µL aliquots, and stored at -80°C until further use.

To prepare protein antigens for coating ELISA plates, genes encoding *H. influenzae* TbpB variants were first cloned (see Supplementary Tables 1 and 2 for additional information) into a T7 expression vector lacking MBP (internal database #: e5632; Supplementary Figure 5) to eliminate the possibility of anti-MBP antibodies present in the mouse serum being detected in the ELISA. The vector also featured a biotin acceptor peptide to allow for *in vivo* biotinylation by the endogenous *E. coli* biotin ligase (7). Following this, the TbpB-encoding vector was transformed into ER2566 cells and the resulting transformants used to inoculate 10 mL of ZYP-5052 autoinduction medium (6) containing 50 µg/mL of ampicillin. The inoculated cultures were then incubated at 37°C for 18 h with shaking. Following this, the cells were harvested by centrifugation at 3,220 x g for 10 min at room temperature, the resulting supernatant decanted, and the pellets re-suspended in Resuspension Buffer. These mixtures were then mixed with glass beads (0.1-mm diameter) in 2-mL Eppendorf tubes and the cells mechanically lysed using a tabletop cell disruptor (“Disruptor Genie,” Scientific Industries). Finally, the resulting lysates were centrifuged at 21,300 x g for 20 min at 4°C and the clarified lysates subsequently diluted and applied directly to streptavidin-coated ELISA wells.
