## Supplementary Tables for "A “loopless” subdomain of transferrin binding protein B can elicit a broadly cross-reactive humoral immune response targeting variants from both encapsulated and non-typeable *Haemophilus influenzae*"

| Primer Name | Primer Sequence | | Anchor Peptide Truncation |
| --- | --- | --- | --- |
| HiV1F11BamHI | | CGCGGATCCGACGTCTCTAATCCCTCCTCTTCTAAA | -11 |
| Hi201F9BamHI | | CGCGGATCCGATAACGTCTCTAATACCCCCTCTTCTAAACC | -9 |
| Hi210Rev-Hind3 | | CCCAAGCTTCTACTTGTTGGTTTTTTCTACTTGTTTTTTAGCTCCAAA | N/A |
| Hi040Rev-XbaI | | ATATTCTAGATTACTTGTTGTTTGTTTCTACTTGTTTTTTAGCTCC | N/A |
| Hi201Rev-XbaI | | ATATTCTAGATTATTTGGTTGTTTCTACTTGTTGTCTCGC | N/A |
| Hi014Rev-XbaI | | ATATTCTAGACTACTTGTTGTTTGTTTCTACTTGTTTTTTAGCTCC | N/A |
| Hi216Rev-Hind3 | | CCCAAGCTTTTACTTGTTGTTTTTTTCTACTTGTTTTTAGCACCAAA | N/A |
| WP_111-TbpB-Rev-HindIII | | CCCAAGCTTTTATTTGGTTGTTTCTACTTGTTTTTTAGCT | N/A |
| WP_11887-TbpB-Rev-HindIII | | CCCAAGCTTTTATTTGGTTGTTTCTACTTGTTTTTTAGCG | N/A |
| WP_221-TbpB-Rev-HindIII | | CCCAAGCTTTTATTTGGTTGTTTCTACTTGTTGTTTTTTAG | N/A |

**Supplementary Table 1:** Primers used to clone *H. influenzae* *tbpB* variants into T7 expression vectors using restriction cloning. Restriction enzyme digestion was conducted using BamHI and either XbaI or HindIII according to the manufacturer’s instructions (ThermoScientific’s FastDigest protocol). The anchor peptide (N-terminal) truncation produced by each forward primer used in this study is indicated where appropriate.

**Supplementary Table 2:** Primer pairs and annealing temperatures used for amplifying each *H. influenzae tbpB* variant examined in this study in preparation for restriction cloning. All PCRs were conducted using the Phire II DNA polymerase (ThermoScientific) according to the manufacturer’s instructions. The sequences for the primers listed here can be found in Supplementary Table 1.

| TbpB Variant | Primer Pair Used for Cloning | | Annealing Temperature  (°C) |
| --- | --- | --- | --- |
| h036 | | Forward: Hi201F9BamHI  Reverse: Hi210Rev-Hind3 | 68.3 |
| h040 | | Forward: HiV1F11BamHI  Reverse: Hi040Rev-XbaI | 67.7 |
| h201 | | Forward: Hi201F9BamHI  Reverse: Hi201Rev-XbaI | 67.1 |
| TbpB-1 (WP_050848381.1) | | Forward: HiV1F11BamHI  Reverse: Hi210Rev-Hind3 | 68.2 |
| TbpB-2 (h014) | | Forward: HiV1F11BamHI  Reverse: Hi014Rev-XbaI | 67.9 |
| TbpB-3 (WP_112083867.1) | | Forward: Hi201F9BamHI  Reverse: Hi040Rev-XbaI | 67.7 |
| TbpB-4 (h026) | | Forward: HiV1F11BamHI  Reverse: Hi201Rev-XbaI | 67.1 |
| TbpB-5 (WP_221267536.1) | | Forward: HiV1F11BamHI  Reverse: WP_221-TbpB-Rev-HindIII | 63.5 |
| TbpB-6 (WP_118806425.1) | | Forward: HiV1F11BamHI  Reverse: Hi210Rev-Hind3 | 68.2 |
| TbpB-7 (PRI69814.1) | | Forward: Hi201F9BamHI  Reverse: Hi210Rev-Hind3 | 68.3 |
| TbpB-8 (PRI75995.1) | | Forward: HiV1F11BamHI  Reverse: Hi216Rev-Hind3 | 68.2 |
| TbpB-9 (h216) | | Forward: HiV1F11BamHI  Reverse: Hi216Rev-Hind3 | 68.2 |
| TbpB-10 (WP_111689186.1) | | Forward: HiV1F11BamHI  Reverse: WP_111-TbpB-Rev-HindIII | 64.2 |
| TbpB-11 (h210) | | Forward: HiV1F11BamHI  Reverse: Hi210Rev-Hind3 | 68.2 |
| TbpB-12 (WP_118872958.1) | | Forward: HiV1F11BamHI  Reverse: WP_11887-TbpB-Rev-HindIII | 64.8 |
