## Supplementary Figures for "A “loopless” subdomain of transferrin binding protein B can elicit a broadly cross-reactive humoral immune response targeting variants from both encapsulated and non-typeable *Haemophilus influenzae*"

### Slide 1
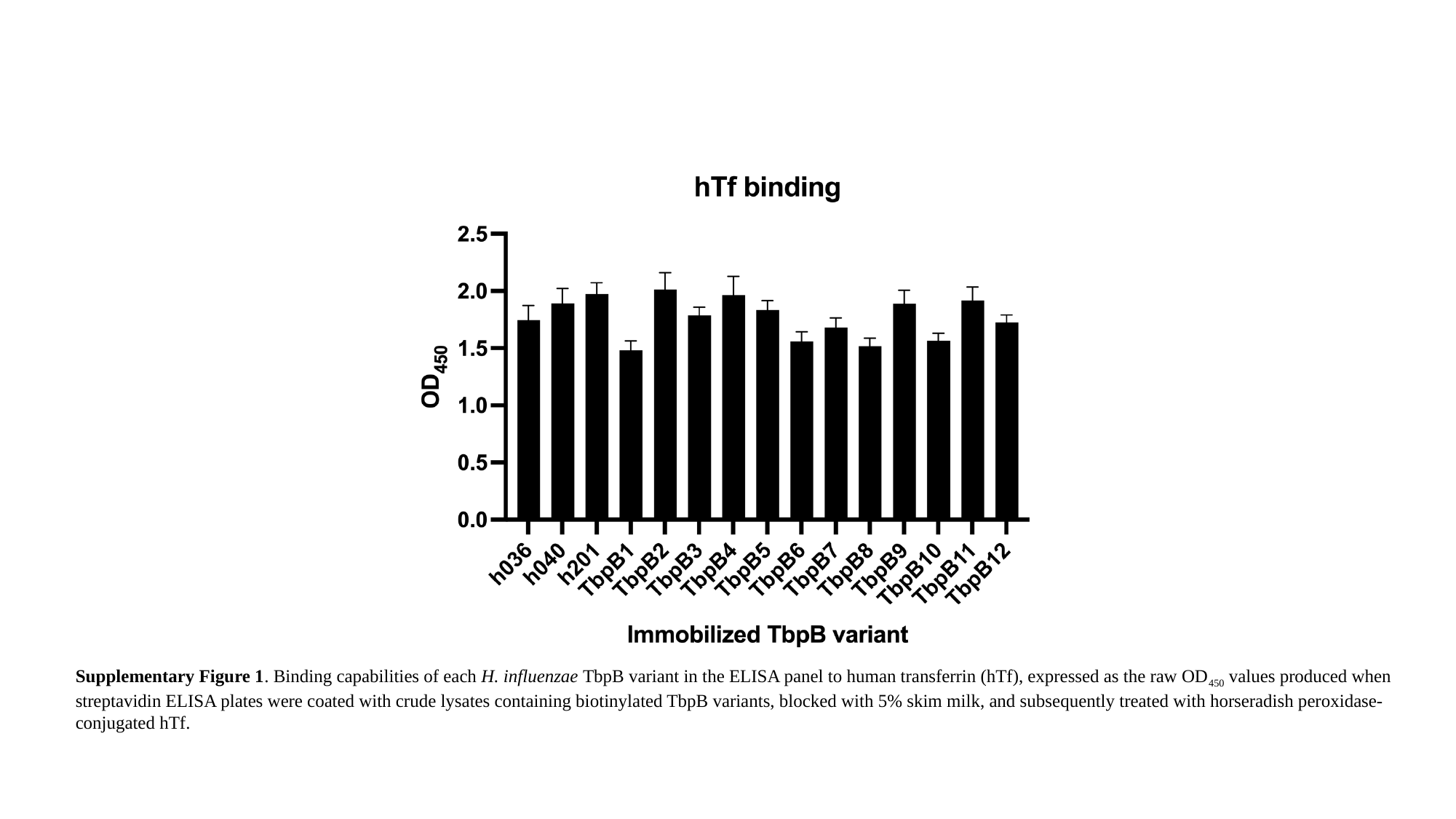

Supplementary Figure 1. Binding capabilities of each H. influenzae TbpB variant in the ELISA panel to human transferrin (hTf), expressed as the raw OD_450_ values produced when streptavidin ELISA plates were coated with crude lysates containing biotinylated TbpB variants, blocked with 5% skim milk, and subsequently treated with horseradish peroxidase-conjugated hTf.

### Slide 2
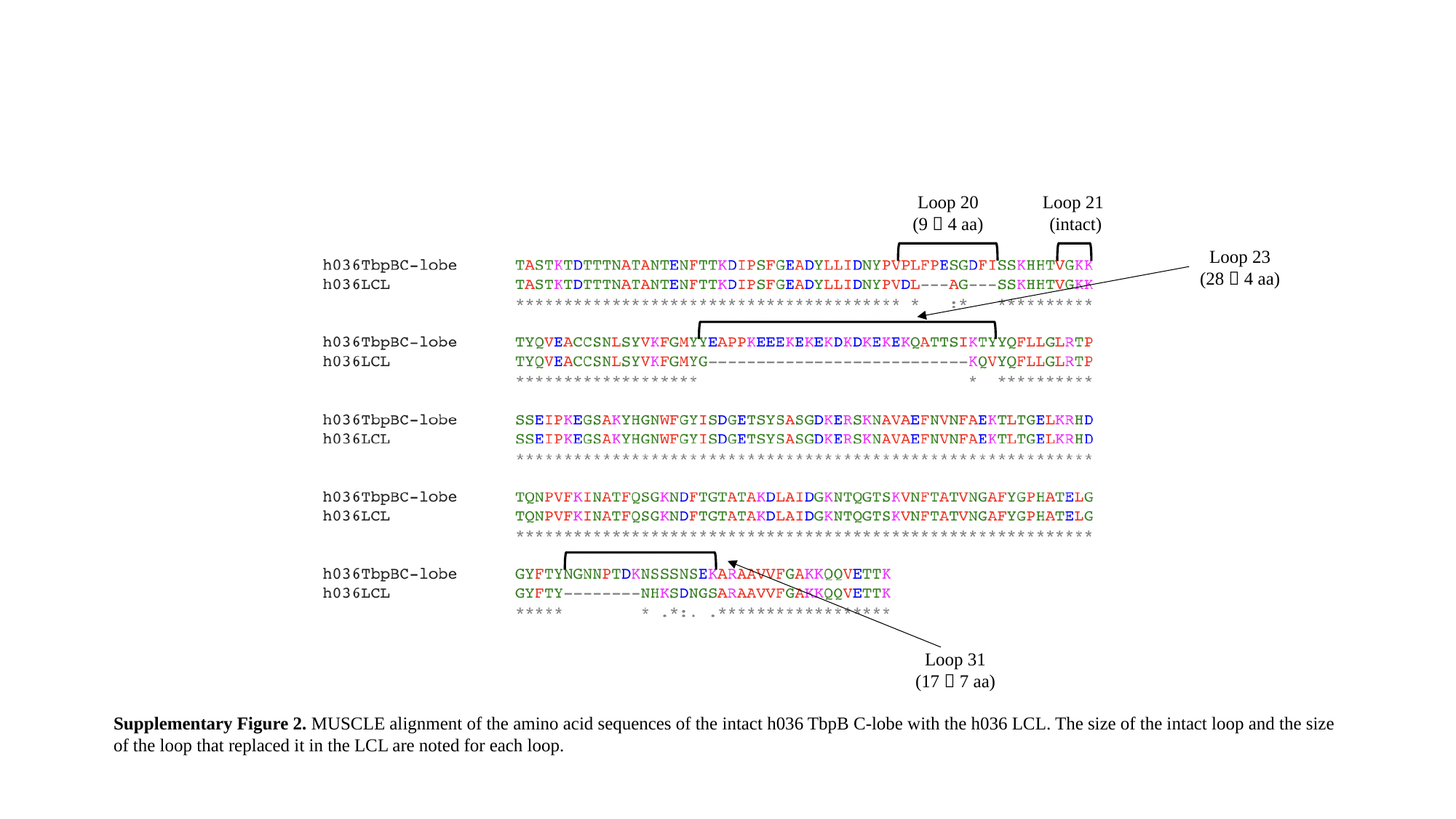

Loop 20
(9 🡪 4 aa)
Loop 21
(intact)
Loop 23
(28 🡪 4 aa)
Loop 31
(17 🡪 7 aa)
Supplementary Figure 2. MUSCLE alignment of the amino acid sequences of the intact h036 TbpB C-lobe with the h036 LCL. The size of the intact loop and the size of the loop that replaced it in the LCL are noted for each loop.

### Slide 3
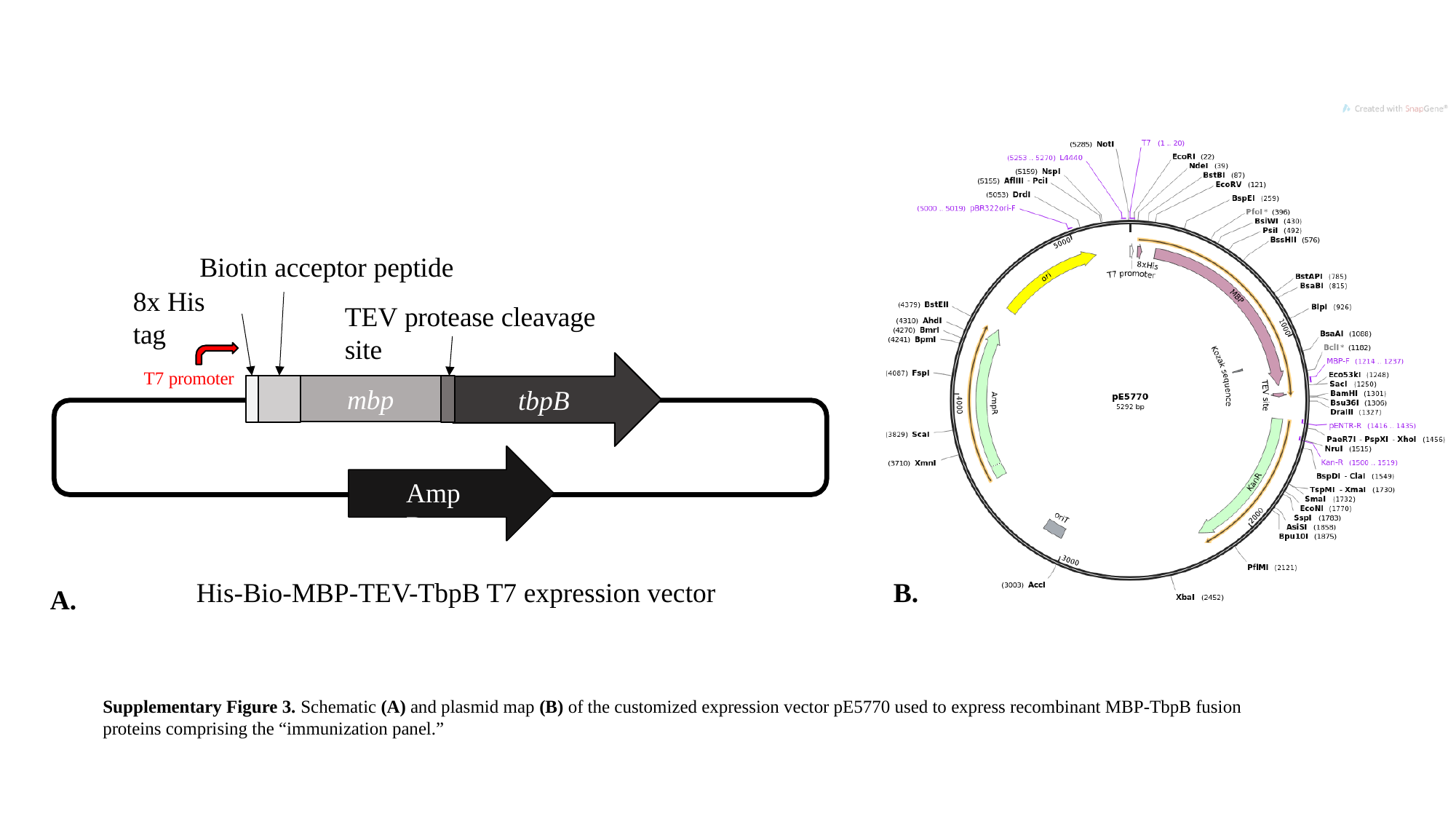

Biotin acceptor peptide
8x His tag
TEV protease cleavage site
T7 promoter
AmpR
tbpB
mbp
His-Bio-MBP-TEV-TbpB T7 expression vector
B.
A.
Supplementary Figure 3. Schematic (A) and plasmid map (B) of the customized expression vector pE5770 used to express recombinant MBP-TbpB fusion proteins comprising the “immunization panel.”

### Slide 4
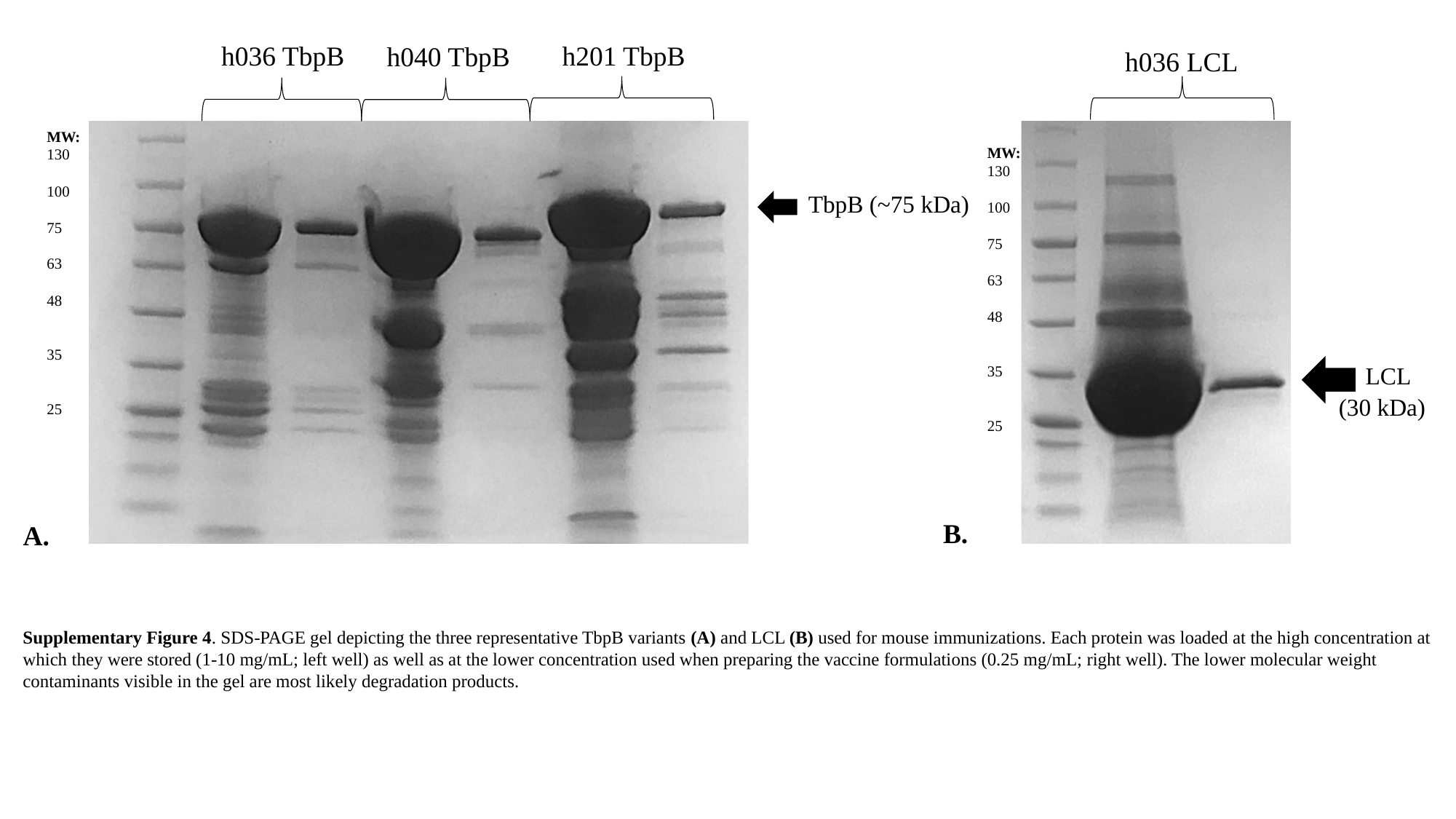

h036 TbpB
h201 TbpB
h040 TbpB
MW:
130
100
75
63
48
35
25
TbpB (~75 kDa)
MW:
130
100
75
63
48
35
25
LCL
(30 kDa)
h036 LCL
B.
A.
Supplementary Figure 4. SDS-PAGE gel depicting the three representative TbpB variants (A) and LCL (B) used for mouse immunizations. Each protein was loaded at the high concentration at which they were stored (1-10 mg/mL; left well) as well as at the lower concentration used when preparing the vaccine formulations (0.25 mg/mL; right well). The lower molecular weight contaminants visible in the gel are most likely degradation products.

### Slide 5
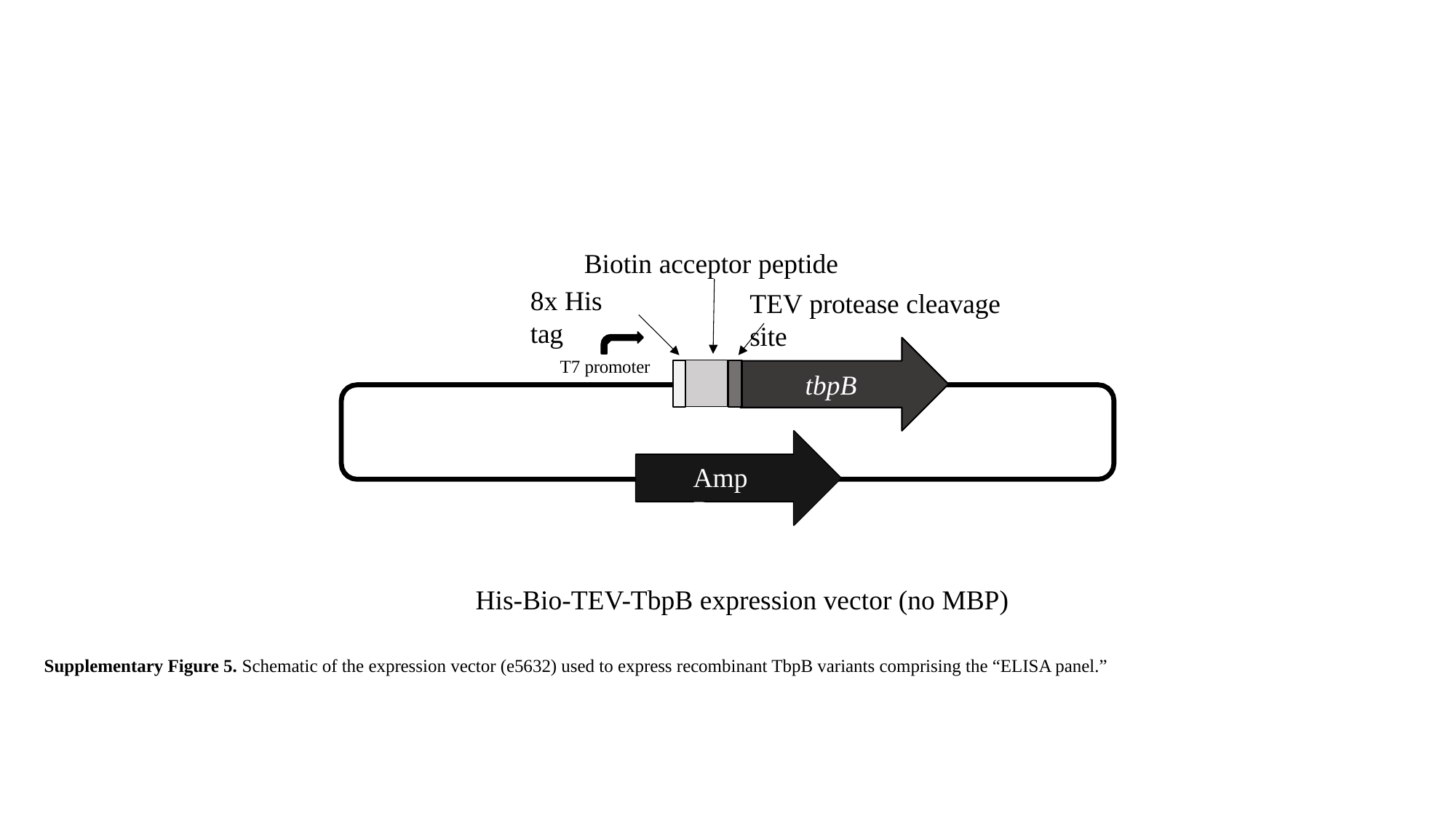

Biotin acceptor peptide
8x His tag
TEV protease cleavage site
T7 promoter
AmpR
tbpB
His-Bio-TEV-TbpB expression vector (no MBP)
Supplementary Figure 5. Schematic of the expression vector (e5632) used to express recombinant TbpB variants comprising the “ELISA panel.”

### Slide 6
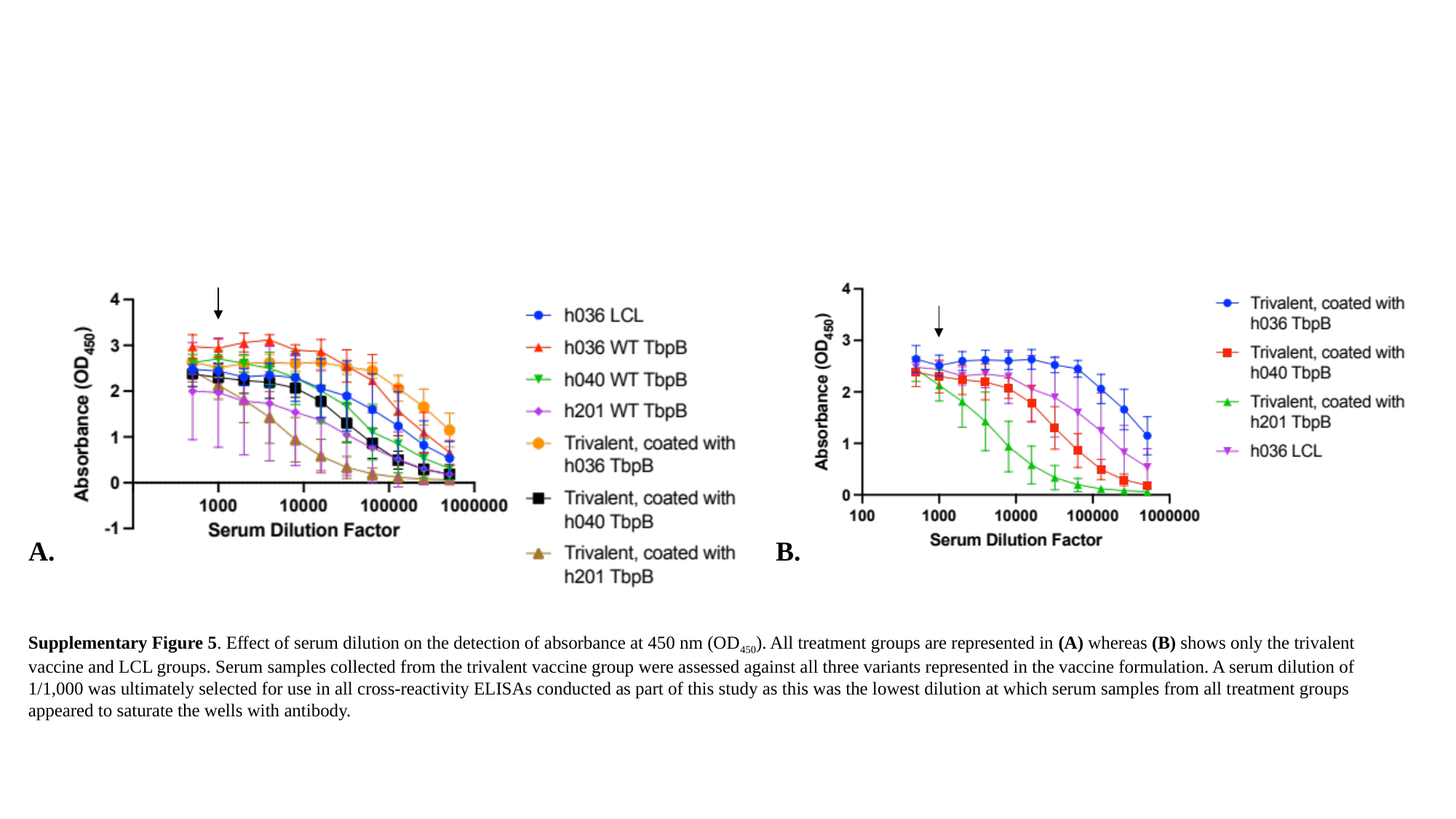

A.
B.
Supplementary Figure 5. Effect of serum dilution on the detection of absorbance at 450 nm (OD_450_). All treatment groups are represented in (A) whereas (B) shows only the trivalent vaccine and LCL groups. Serum samples collected from the trivalent vaccine group were assessed against all three variants represented in the vaccine formulation. A serum dilution of 1/1,000 was ultimately selected for use in all cross-reactivity ELISAs conducted as part of this study as this was the lowest dilution at which serum samples from all treatment groups appeared to saturate the wells with antibody.
